# A Negative Feedback Regulates The Flow Of Signal Through Akt/mTORC1/S6K1 Pathway

**DOI:** 10.1101/147710

**Authors:** Poulami Dutta, Vimalathithan Devaraj, Biplab Bose

## Abstract

Several growth factors, cytokines, hormones activate PI3K/Akt pathway. Akt is a key node in this pathway and activates different downstream paths. One such path is Akt/mTORC1/S6K1 that controls protein synthesis, cell survival, and proliferation. Here we show that a negative feedback controls activation of S6K1 through this pathway. Due to this negative feedback, a sustained phospho-Akt signal generates a transient pulse of phospho-S6K1. We have created a mathematical model for this circuit. Analysis of this model shows that the negative feedback acts as a filter and preferentially allows a signal, with sharp and faster rise of phospho-Akt, to induce pronounced activation of S6K1. It blocks an input with a fast oscillation of phospho-Akt to flow through this path. We show that this negative feedback leads to differential activation of S6K1 by Insulin and Insulin-like Growth Factor 1. Such differential effect may explain the difference in the mitogenic effect of these two molecules.

## Introduction

Signaling molecules, like hormones, growth factors, bind to specific cell surface receptors and activate one or more signaling pathways. Though receptors have high specificity, a cell has only a few canonical pathways to process external signals. This gives rise to the ‘hourglass’ structure of signaling network, where signals generated by different signaling molecules are processed by a handful of core pathways (1–2). Eventually, the hourglass widens again as signals propagated through core pathways lead to diverse cellular functions.

Signaling pathways involve cascades of enzymatic reactions that covalently modify proteins and change their activity or stability. The concentrations, modification states, and activities of pathway molecules change with time. This temporal dynamics of pathway molecules carry the message given by an input signal and decide cellular response (3–5). Both identity and strength of the external signal affect this temporal dynamics (4). Temporal dynamics of pathway molecules also depend on the architecture of a pathway (4).

The network architecture also decides the path through which a particular input signal can propagate effectively. Two input signals may activate same upstream molecules in a pathway, but eventually may pass through different downstream branches, giving rise to different cellular responses. Such discrimination depends upon network motifs involved in the pathway (6).

PI3K/Akt pathway is a cardinal pathway for cell signaling. A large number of signaling molecules activate this pathway (7). It controls diverse cellular processes like cell survival, proliferation, glucose metabolism, and cell motility (8–10). Akt is a key node in this pathway. Activation of PI3K/Akt pathway triggers phosphorylation of Akt. Akt has multiple substrates, and from Akt, the pathway splits into multiple branches (8,11). However, Akt does not activate all the downstream branches equally. There exists considerable heterogeneity in Akt-mediated activation of its substrates (12).

One such branch, from Akt, involves mTORC1. Signaling through mTORC1 regulates translational machinery and is crucial in the regulation of cell size, cell cycle, and survival (13). Akt indirectly activates mTORC1, and in turn, mTORC1 activates p70-S6K1 (S6K1) through phosphorylation (13–14). Active S6K1 regulates several molecules involved in protein synthesis, namely RPS6, PDCD4, and EIF4B (15). It is also involved in processes linked to cell survival, proliferation, cytoskeletal rearrangement, and metabolism (16–17).

Recently, Rahman and Haugh (18) proposed that Akt controls activation of mTORC1 by a feedforward motif and S6K1 is linearly controlled by mTORC1. Following this network architecture, the temporal behavior of phospho-S6K1 follows the temporal behavior of phospho-Akt. However, Kubota *et al.* (19) had earlier proposed that an incoherent feedforward (IFF) controls S6K1 in Akt/mTORC1/S6K1 pathway. They proposed this network motif to explain the transient behavior of phospho-S6K1 observed in their experiments.

In the present work, we investigate the signal transfer from Akt to S6K1. We show that phosphorylation of S6K1 is regulated by a negative feedback, not an IFF. The negative feedback works as an adaptive motif and transforms a sustained signal by phospho-Akt (pAkt) into a transient phospho-S6K1 (pS6K1) response. Further, we have developed a mathematical model to understand the transformation of temporal dynamics of pAkt into the temporal dynamics of pS6K1. Temporal dynamics of pAkt varies with the dose and type of external cue. The negative feedback ensures that only certain input signals would be able to activate the Akt/mTORC1/S6K1 pathway effectively.

## Results

### Temporal dynamics of phosphorylation of Akt and S6K1

Insulin-like growth factor 1 (IGF-1) binds to its receptor on a cell and activates PI3K/Akt pathway (20). We have treated MCF-7 cells with 5 nM and 10 nM of IGF-1 and detected phosphorylation of Akt, at different time points. In both experiments, phospho-Akt (pAkt) increased with time eventually reaching a steady state (Figure 1). The rise and the steady state of pAkt varied with the dose of IGF-1. The temporal dynamics of pAkt fitted well with the Hill function 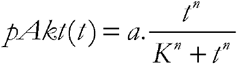. The amplitude (a), Hill constant (K) and Hill coefficient (n) depend upon the dose of IGF-1 used. This indicates that the dose-dependent message, provided by IGF-1, is coded in the temporal dynamics of pAkt.

**Figure 1:**
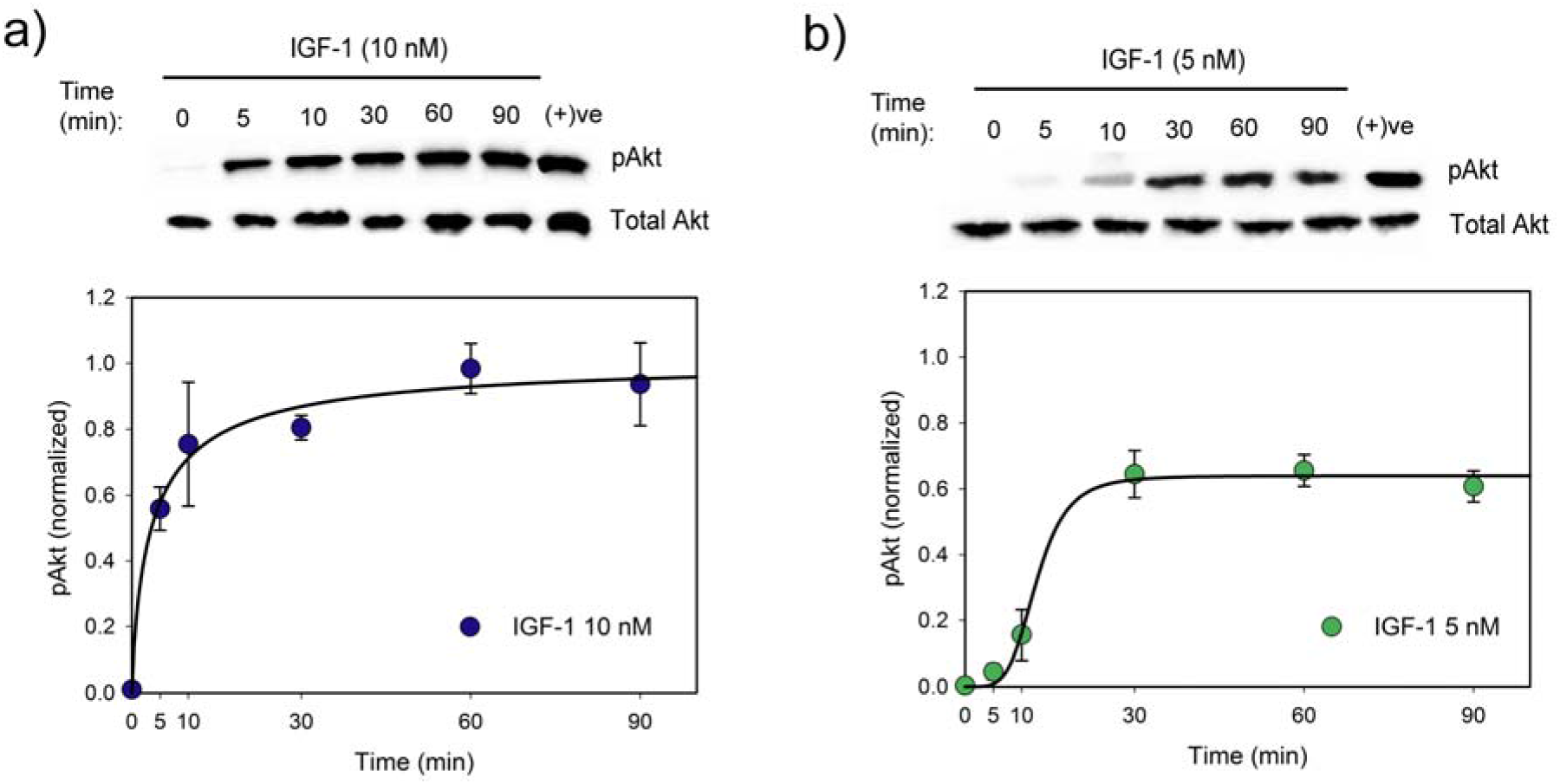
Temporal dynamics of phosphorylation of Akt. MCF-7 cells were treated with (a) 10 nM and (b) 5 nM of IGF-1 and phospho-Akt (Ser473) was detected by Western Blot. Representative blots are shown here. Quantitative data from densitometry are shown in the graphs below respective blots. Each data point represents mean of three independent experiments, and error bars indicate standard deviation. Data points are fitted to Hill functions. For 10 nM IGF-1, 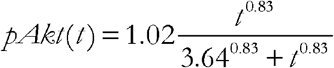. For 5 nM IGF-1, 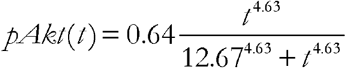. (+)ve: cells maintained in media with 10% serum.

This coded message should be decoded at the lower level of the pathway. IGF-1 activates PI3K/Akt pathway. Akt activates mTORC1 which in turn phosphorylates S6K1 (14). We measured the temporal dynamics of phospho-S6K1 (pS6K1) in IGF-1-treated cells.

When treated with IGF-1, pS6K1 showed a transient response (Figure 2a and b). It had an initial rise, with an eventual decline. By 90 minutes, pS6K1 returned to a lower level, even though pAkt remained at the higher level. This behavior was observed for both the doses of IGF-1. However, the amplitude of the transient rise of pS6K1 varied with the dose of IGF-1.

**Figure 2:**
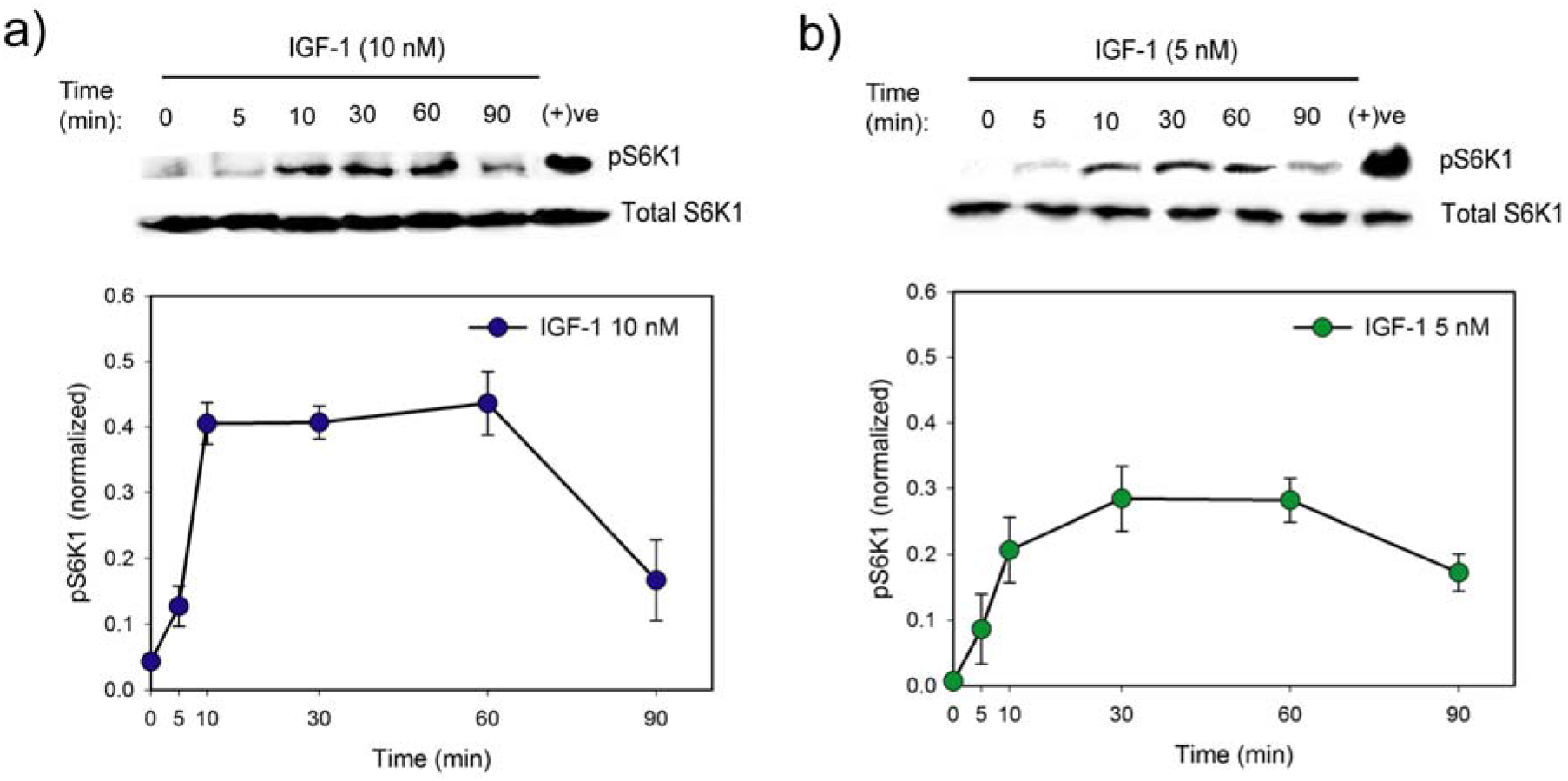
Temporal dynamics of phosphorylation of S6K1. MCF-7 cells were treated with (a) 10 nM and (b) 5 nM of IGF-1 and phospho-S6K1 (T389) was detected by Western Blot. Representative blots are shown here. Quantitative data from densitometry are shown in the graphs below respective blots. Each data point represents mean of three independent experiments and error bars indicate standard deviation. (+)ve: cells maintained in media with 10% serum.

We did experiments to confirm that IGF-1 is inducing phosphorylation of S6K1, only, through PI3K/Akt/mTORC1 pathway. Cells were treated with IGF-1 in the presence of PI3K inhibitor (LY294002) and mTORC1 inhibitor (Rapamycin). PI3K inhibitor blocked IGF-1-induced phosphorylation of Akt, and S6K1 (Supplementary figure S1a). On the other hand, Rapamycin blocked phosphorylation of S6K1 without affecting phosphorylation of Akt (Supplementary figure S1b).

We treated MCF-7 cells with IGF-1 in the presence of U0126, an MEK1/2 inhibitor. However, U0126 had no effect on the IGF-1-induced temporal dynamics of pAkt, and pS6K1 (Supplementary figure S1c). This confirmed that IGF-1 induced phosphorylation of S6K1 does not involve MAPK pathway in these cells. DMSO was used as the vehicle for all the pathway inhibitors used in this work. It does not affect phosphorylation of Akt and S6K1 (Supplementary figure S1d).

### Network motif that controls phosphorylation of S6K1

The transient time course of pS6K1 is typical of an adaptive network motif (21–22). In an adaptive motif, sustained input gives rise to a transient pulse of output. Both negative feedback and incoherent feedforward cause such transient behavior (22). Figure 3a shows three possible network motifs that can give rise to the observed temporal dynamics of pS6K1. First two motifs (I and II) are negative feedbacks. The third motif is an incoherent feedforward (IFF). Ma *et al.* (22) have shown that motif II and III show robust adaptive behavior for a broad range of parameter values.

**Figure 3:**
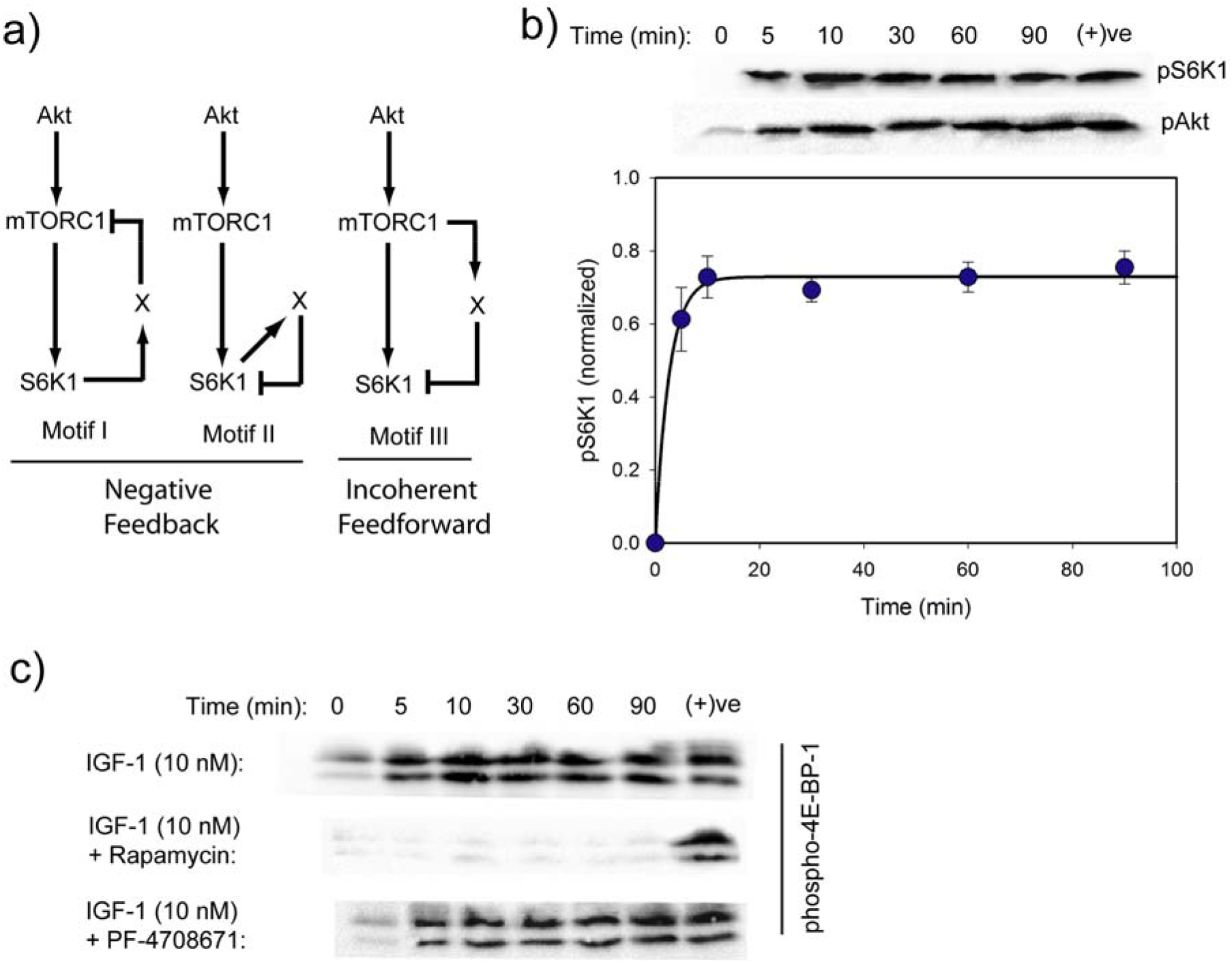
Network motif that controls S6K1. a) Three possible network motifs that will give rise to the adaptive behavior observed for phospho-S6K1. b) Time-dependent Western blot for phospho-S6K1 in MCF-7 cells treated with IGF-1 (10 nM) in the presence of S6K1 inhibitor PF-4708671. A representative blot is shown here with quantitative data from three independent experiments presented in the graph below the blot. Error bars indicate standard deviation. c) Time-dependent Western blots to detect phospho-4E-BP-1 (Thr37/46) in MCF-7 cells treated with IGF-1 (10 nM) alone and in the presence of Rapamycin and PF-4708671. (+)ve: cells maintained in media with 10% serum.

We performed experiments to discriminate between these three motifs. PF-4708671 is a specific inhibitor of the kinase activity of S6K1. It does not affect the basal level of phosphorylation of Akt and S6K1 in MCF-7 cells (Supplementary figure S2). However, inhibition of S6K1 activity, by PF-4708671, would remove the negative feedback in motif I and II. This will linearize these two motifs, and the adaptive property of these motifs would be lost. Therefore, cells co-treated with IGF-1 and PF-4708671 would show sustained activation of S6K1. However, inhibition of S6K1 activity will not affect motif III.

We treated MCF-7 cells with IGF-1 in the presence of PF-4708671. We observed that in the presence of this inhibitor, the transient behavior of pS6K1 is lost (Figure 3b). Like pAkt, pS6K1 also increased with time and reached a higher steady state. This loss of adaptive behavior rules out motif III, the IFF.

Apart from S6K1, mTORC1 phosphorylates several other molecules (23). In motif I, the negative feedback from S6K1 inhibits mTORC1 activity. That would affect the dynamics of phosphorylation of any substrate of mTORC1. Inhibition of S6K1 would also change the dynamics of phosphorylation of those substrates. We explored this feature to discriminate between motif I and II.

mTORC1 phosphorylates 4E-BP-1 (24). We treated MCF-7 cells in presence/absence of different inhibitors and observed the temporal dynamics of phospho-4E-BP-1 by Western Blot. We observed that IGF-1 induces rapid phosphorylation of 4E-BP-1 and phosphorylation reaches the steady state at an early time point (Figure 3c upper panel). Treatment with Rapamycin inhibits such phosphorylation (Figure 3c middle panel). This confirms that mTORC1 phosphorylates 4E-BP-1. However, inhibition of S6K1 did not affect the temporal dynamics of phsopho-4E-BP-1 (Figure 3c lower panel). These observations indicate that the negative feedback from S6K1 does not affect mTORC1 activity and rule out motif I.

### A mathematical model for the network motif of S6K1

Our experiments confirmed that a negative feedback, similar to motif II of Figure 3a, controls phosphorylation of S6K1. We created a mathematical model for this network motif, using ordinary differential equations. The parameters of the model were estimated from the results of experiments, where cells were treated with IGF-1 (5 and 10 nM) in presence and absence of S6K1 inhibitor. The details of the model are provided in the supplementary text. Akt activates mTORC1 through multiple steps, involving TSC1/2, Rheb-GTP, and PRAS40 (13-14,18). However, this part of the pathway, from Akt to mTORC1, does not involve any feedback or incoherent feedforward. Our results also show that in MCF-7 cells inhibition of mTORC1 by Rapamycin does not affect the temporal dynamics of pAkt (Supplementary figure S1b). Therefore, signal transfer from Akt to mTORC1 is not responsible for the observed adaptive behavior of S6K1 and possibly only rescaling the signal from pAkt. Considering these, we have removed mTORC1 from our model and connected Akt directly to S6K1. This makes the model simpler but still allows us to explore the behavior of the negative feedback, as motif II does not involve any molecule upstream to S6K1. The architecture of the model is shown in Figure 4a.

**Figure 4:**
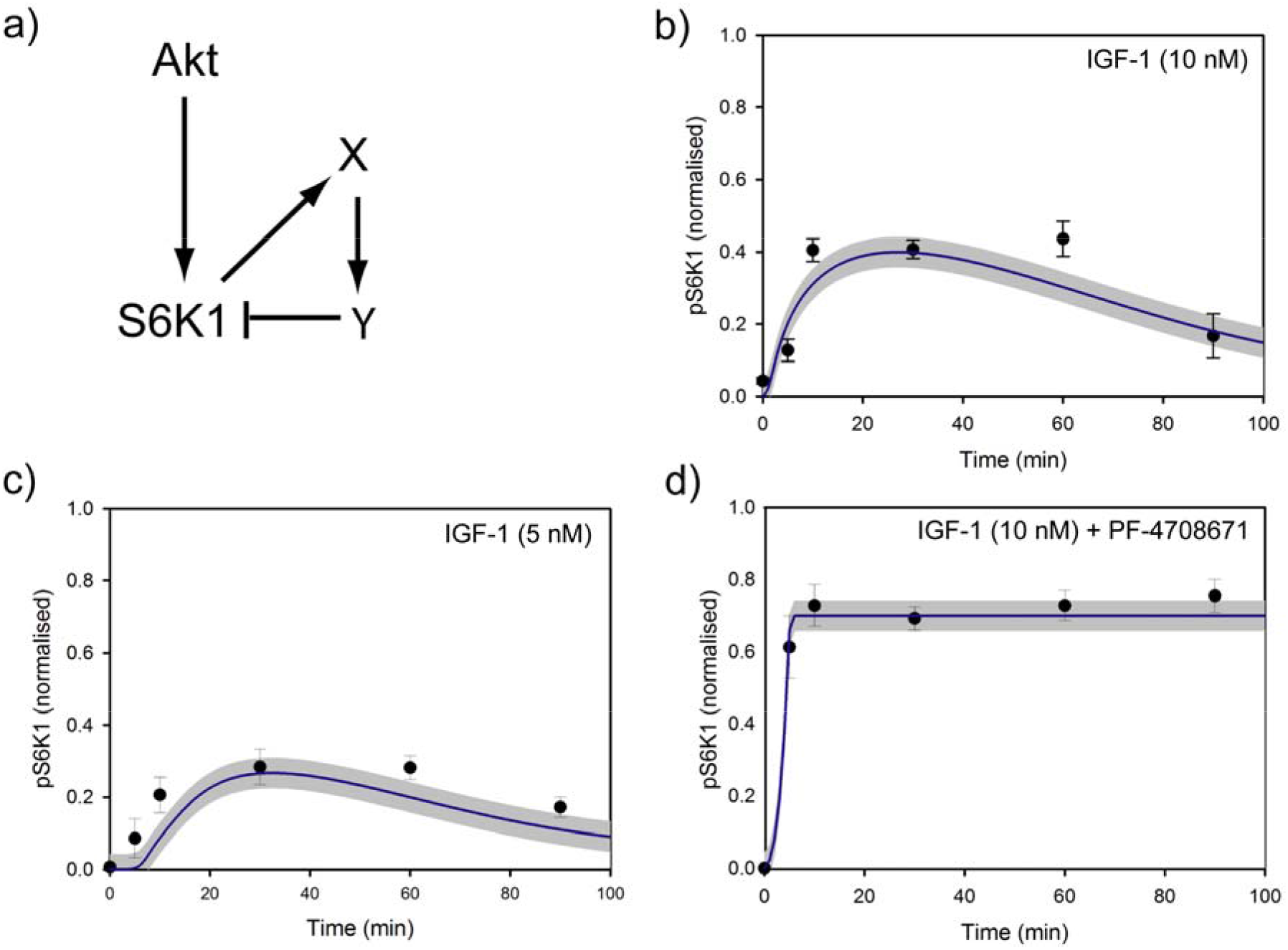
Modeling the network motif for S6K1. a) The architecture of the molecular circuit used to model the negative feedback network motif for S6K1. b) to d) Fitting of model outputs with the experimental data that has been used for parameter estimation. Treatment conditions are written over the plots. Filled circles with error bar are experimental results. Solid lines represent the output of the model. Grey region around solid line represents the estimated error.

We have considered two buffer molecules X and Y in the negative feedback (Figure 4a). Buffers in negative feedback introduce delay. Delay in a negative feedback is necessary to produce a transient adaptive output (21,25). The transient time course of pS6K1 was not a sharp pulse, but a broader one. Two buffer molecules, rather than one, in the negative feedback, allowed us to fit our model better to experimental data. Fitting of our model outputs to experimental results is shown in Figure 4 (b-d).

To validate the mathematical model, we have performed additional experiments. In one, MCF-7 cells were treated with 10 nM Insulin and time course of phosphorylation of Akt and S6K1 was measured. Insulin also activates PI3K/Akt pathway. In another experiment, cells were treated with both Insulin and IGF-1, but at lower doses (2.5 nM each). The data of these experiments are shown in Figure 5a and b (upper panels). The temporal dynamics of pAkt in both the experiments fitted well with Hill functions (Figure 5 a and b, graphs in the middle). These Hill functions were used as input for our model and dynamics of pS6K1 were predicted by simulation. Simulated data matched reasonably well with the observed behavior of pS6K1 (Figure 5 a and b, graphs in lower panel).

**Figure 5:**
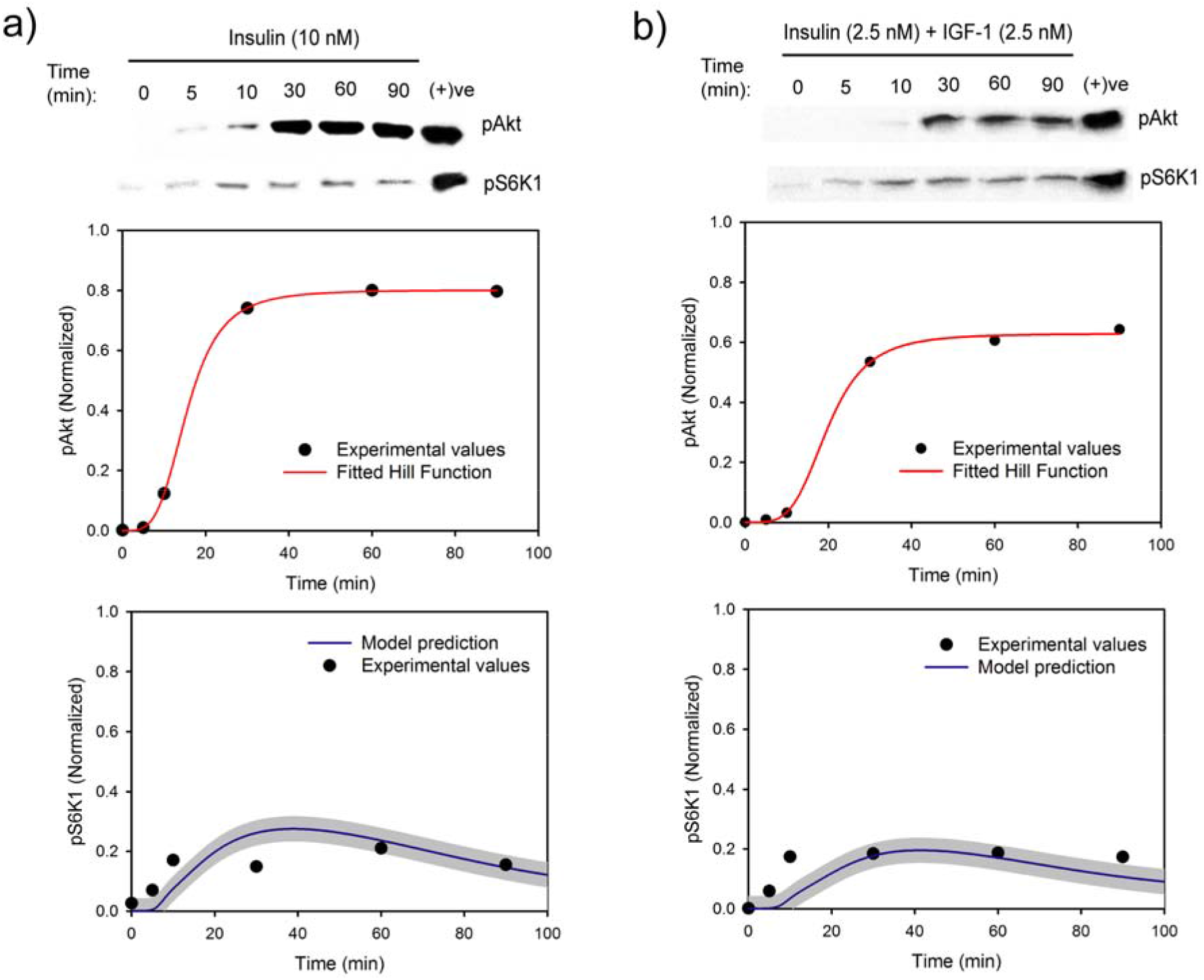
Validation of the mathematical model. Cells were treated with either (a) 10 nM Insulin or (b) Insulin and IGF-1 (2.5 nM each). Levels of pAkt and pS6K1 at different time points were measured by Western blot (upper panels) and quantified by densitometry. Time course data of pAkt in both the experiments (solid circles, middle graphs) were fitted to Hill functions (Red lines). Lower graphs show the quantified time course of pS6K1 in both the experiments (solid circles). Temporal behaviors of pS6K1, as predicted by the model, are shown by blue lines (lower graphs). Shaded regions represent the estimated error. (+)ve: cells maintained in media with 10% serum.

### The negative feedback circuit as signal decoder

Our experiments highlight a general characteristic of molecular signaling. Both insulin and IGF-1 activates PI3K/Akt pathway. However, the temporal dynamics of pAkt differs with ligand and dose of the ligand used. In other words, the message given by a specific dose of a particular ligand is encoded in the temporal dynamics of pAkt. The dynamics of pAkt is transformed into specific temporal dynamics of pS6K1, through the negative feedback circuit. Our experiment and mathematical model show that the dynamics of pS6K1 depends solely on the time course of pAkt, irrespective of the ligand used to activate the pathway.

One can consider the negative feedback motif for S6K1 as a decoder that decodes the message encoded in the time course of pAkt. To work as a decoder, this circuit involving S6K1 should be able to differentiate different temporal dynamics of pAkt. Both transient and sustained temporal dynamics of pAkt are reported in the literature (19,26-28). We have used two different temporal dynamics of pAkt as input and simulated the behavior of pS6K1. These are: a) pAkt follows a sigmoidal time course, as observed in our experiments and b) A transient pulse of pAkt.

For all these inputs, pS6K1 has a transient response, with an initial rise and subsequent decay to a lower steady state. Although the steady state values changed with input signals, those are very close to each other. Therefore, we characterized the temporal dynamics of pS6K1 in terms of the amplitude (height of the peak) and the area under the curve (AUC) of pS6K1 vs. time plot. The area under the curve (AUC) represents time-integral of cumulative activation of S6K1.

For sigmoidal inputs, we have used Hill functions with constant amplitude but different Hill coefficient and Hill constants. The Hill constant decides the delay in the rise of pAkt On the other hand, the Hill coefficient determines how sharp or fast it increases. Both the amplitude and AUC of pS6K1 decrease with increase in Hill constant (Figure 6a and b). That means the negative feedback circuit of S6K1 can differentiate an input signal that rises early from a delayed one. Both AUC and amplitude of pS6K1 are high for higher values of Hill coefficient (6a and b). Therefore, this motif can also differentiate between a slow and a fast input.

**Figure 6.**
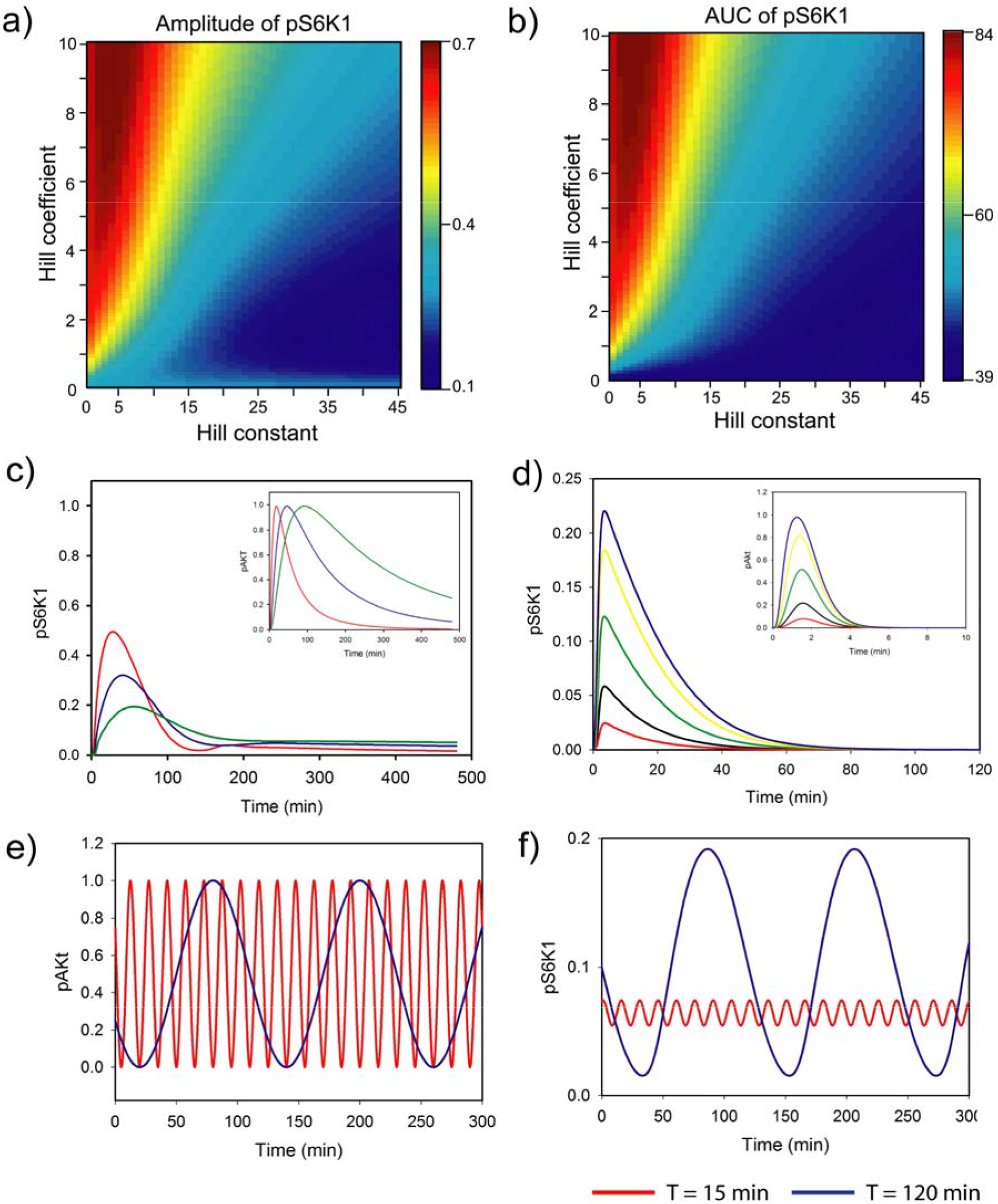
Input-output characteristics of the negative feedback. Different temporal dynamics of pAkt were used as inputs and temporal behavior of pS6K1 was estimated by simulation. a) and b) show the behavior of pS6K1 for sigmoidal inputs. Hill functions with different Hill coefficients and Hill constants were used. The amplitude of pAkt is kept constant at 1. c) and d) show the behavior of pS6K1 for transient inputs as shown in corresponding insets. Color codes in main graphs are same as in corresponding inset plots. e) Shows two oscillating input signals with different periods (T). f) Temporal dynamics of pS6K1 for oscillatory input signals. The stable trajectories of pS6K1 are shown. The color code is same as in (e).

Negative feedback is an adaptive motif. The behavior of pS6K1 depends upon how fast the input increases and the adaptation time scale. With delayed, slow rising input, the system adapts quicker than the rise in input and thereby does not generate pronounced output. Therefore, this motif facilitates a selective transfer of fast rising input signal to downstream molecules.

This property is also observed for transient input signals. We performed simulations with three different pulses of pAkt as input. All of these have equal amplitude but have different slopes for the rise (inset of Figure 6c). As shown in Figure 6c, the amplitude of pS6K1 depends upon the rate of rise of pAkt and a sharp rise in pAkt causes higher amplitude for pS6K1. In another simulation, we have used pulses of pAkt having different amplitudes (inset of Figure 6d). It was observed that the amplitude of pS6K1 increases with increase in the amplitude of pAkt (Figure 6d).

### The negative feedback circuit filters out input with fast oscillation

In some cases, activation of PI3K/Akt pathway generates oscillation of pAkt (29–30). Sometimes the input signal itself is oscillatory. For example, insulin level in human plasma has temporal oscillation (31). Two different pulses of Insulin are usually observed, one with shorter a period (~ 12 minutes) and another with a longer period (1- 3 hr) (32–33). Such oscillatory input would generate oscillation of pAkt (19).

We have performed simulations to understand how the negative feedback motif of S6K1 handles such oscillatory input. Two oscillatory signals were used as inputs for these simulations, one with the shorter period (15 minutes) and the other with a period of 120 minutes (Figure 6e). Amplitudes of both the inputs were same. For such oscillatory inputs, pS6K1 shows limit-cycle oscillation. Figure 6f shows the oscillation of pS6K1 in respective stable trajectories. It shows that pS6K1 responds preferentially to a pAkt oscillation with longer period and has oscillation with much higher amplitude. Therefore, this negative feedback filters out an input with faster oscillation in pAkt.

Temporal dynamics of buffers X and Y explains this phenomenon. For an adaptive motif, recovery time is an important parameter. It is the time required for the system to reset itself to the initial state once the input signal is removed (25). An adaptive motif can not respond to successive pulses of input signals, if the negative regulators do not go back to the initial state, after the decay of the first input pulse. Therefore, the period of oscillation of the input signal has to match with the recovery time of the negative feedback motif. In our system, a period of 15 minutes, for pAkt oscillation, is too short with respect to the time required for X and Y to return to basal levels. Therefore, this input can not produce pronounced oscillation in pS6K1.

### Effect of differential activation of S6K1 by IGF-1 and Insulin

We have observed that equal amount of IGF-1 and Insulin differentially activates Akt (Figure 1a and 5a). 10 nM of IGF-1 induced a rapid increase in pAkt; whereas 10 nM of Insulin triggered a delayed rise in pAkt. However, for both IGF-1 and Insulin, steady state values of pAkt were similar. The negative feedback of S6K1 transforms this difference in pAkt dynamics into a marked difference in phosphorylation of S6K1 (Figure 2a and Figure 5a). IGF-1 induced a transient rise in pS6K1 well above its basal level. However, for Insulin, it was very close to the basal level.

We investigated the cellular significance of the difference in temporal dynamics of pS6K1 in IGF-1 and Insulin treated cells. S6K1 is one of the key molecules through which PI3K/Akt pathway controls cell survival and proliferation. We treated MCF-7 cells with different doses of Insulin and IGF-1 and measured the change in cell number with respect to untreated cells (Figure 7a). We observed that the cell number increased even at a lower dose of IGF-1 (5 nM). On the other hand, Insulin does not have such pronounced effect on cell number even at the highest dose (25 nM).

Further, we explored the correlation between temporal dynamics of pS6K1 and fold change in cell number. We measured the time-dependent change in pAkt for cells treated with three doses of IGF-1 and Insulin (5, 10, and 25 nM) by Western Blot. The data were fitted to Hill equations and used as inputs for the mathematical model to simulate the temporal dynamics of pS6K1. We estimated the amplitude of pS6K1 for each case from the simulated data. We observed that the amplitude of pS6K1 has reasonable correlation with the fold change in cell number (Figure 7b). S6K1 is not the sole regulator of cell proliferation and survival. Even then, the difference in its activation by IGF-1 and Insulin apparently correlates with the difference in their effect on cell number.

**Figure 7.**
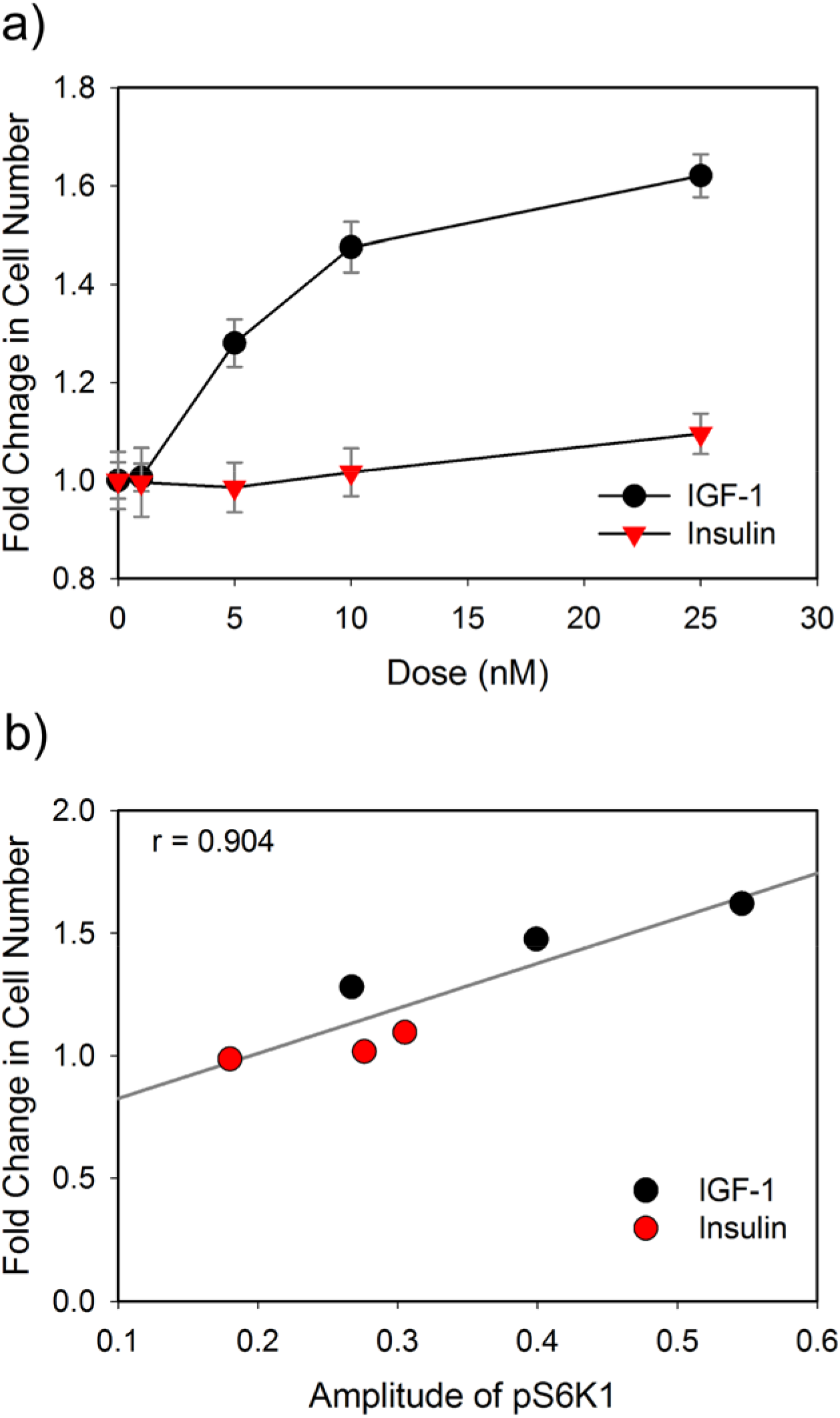
Correlation between S6K1 signaling and fold change in cell number. a) Fold change in cell number at different doses of IGF-1 and Insulin. b) Correlation between fold change in cell number and temporal dynamics of pS6K1 in terms of the amplitude of pS6K1. The amplitude of pS6K1 was estimated from the simulated data. r = Pearson Product-moment Correlation Coefficient.

## Discussion

In this work, we have shown that a negative feedback controls phosphorylation of S6K1 at T389. Phosphorylation at this site is a marker for the active form of S6K1 (34). Due to this negative feedback, a sustained pAkt signal generates only a transient pulse of pS6K1. We have created a mathematical model for this circuit. The model was analyzed to understand the properties of this negative feedback. This negative feedback acts as a filter in the PI3K/Akt/mTORC1/S6K1 pathway and preferentially allows a signal, with sharp and faster rise of pAkt, to induce pronounced activation of S6K1. It does not permit a pAkt oscillation with a shorter period to produce a considerable effect on pS6K1. A large number of signaling molecules activates PI3K/Akt pathway. We have used IGF-1 and Insulin in our experiments. It was observed that the temporal dynamics of pAkt depends upon the signaling molecule and its dose. For both IGF-1 and Insulin, pAkt had sustained response in MCF-7 cells. Such sustained activation of Akt has been observed earlier in MCF-7 cells and some other cell lines (12,28).

However, pS6K1 showed a transient response. Transient increase of pS6K1 has been observed earlier in several other experimental systems (19,35–36). In our experiments, pS6K1 had a transient rise, even though pAkt persistently remained at a higher level. Such adaptive dynamics of pS6K1 has also been reported earlier (12).

This transient pulse of pS6K1 indicates the existence of an adaptive motif that controls phosphorylation of S6K1. Negative feedback and incoherent feed-forward circuits are adaptive motifs that can generate transient outputs (22). Kubota *et al*. (19) had earlier proposed that an incoherent feedforward (IFF) control phosphorylation of S6K1. However, by experiments using inhibitors we have shown that a negative feedback, not an IFF, controls phosphorylation of S6K1.

We have created a mathematical model for this network motif. This model predicted the behavior of pS6K1 when different inputs were used to activate PI3K/Akt pathway. The parameter values of this model have been estimated from our experiments using MCF-7 cells. Therefore, this model may not be suitable for making exact quantitative predictions for other cellular systems. However, our model can be used to provide a qualitative explanation for the transient response of S6K1 observed in other cellular systems (like in (12,19)).

IRS-1, a molecule upstream to Akt in PI3K/Akt pathway, is modulated by a negative feedback from S6K1 (37–38). The temporal dynamics of pAkt is the input signal for our model and inhibition of S6K1 did not have a considerable effect on pAkt dynamics in our experiments (Supplementary Figure S3). Therefore, we have not considered any feedback from S6K1 to upstream of pAkt, in our model.

A delay between activation by the input and the inhibition by the feedback is required to produce a transient output in a negative feedback circuit (25). We have considered two buffer molecules, X and Y, in our model to match the transient behavior of pS6K1 observed in experiments. The identities of X and Y are not yet known. They may be molecules that selectively modulate association of S6K1 with mTOR. Very recently, Liu *et al*. (39) have shown that LY-2779964, an inhibitor of S6K1, increases association of S6K1 with mTOR. Translocation of molecules in different cellular compartments can also work as buffers and introduce a time delay in a pathway (40–41). S6K1 is located predominantly in the cytoplasm. However, growth factor-induced phosphorylation at T389 causes its translocation to the nucleus (42–43). Such translocation of the phosphorylated and active form of S6K1 may cause a delay in the negative feedback.

Due to the negative feedback, a high and persistent input signal, gives rise to a transient increase in pS6K1 and after some time the level of pS6K1 goes back to a lower steady state level. Unless the input is removed and the system is reset, any additional input would not be able to increase pS6K1. We have observed this phenomenon by repeated treatment with high dose of IGF-1 (Supplementary Figure S4). This assures that a high input signal can pass only transiently through the mTORC1/S6K1. However, any aberration in expression or activity of components involved in this negative feedback would disturb this transient behavior. When these molecules are absent or expressed at a lower level, even a weak but persistent input signal will generate a sustained high level of pS6K1 (Supplementary Figure S5). This will change the downstream processes controlled by S6K1.

One can assume that such an aberration may be involved in certain diseases. mTORC1/S6K1 is involved in nutrient sensing, obesity, and Insulin resistance (44). Several studies have shown that mice fed with high-fat diet had sustained elevated phosphorylation of S6K1 in the liver and islets cells than those fed with normal food (45-46). Similar persistent high pS6K1 was observed in db/db and ob/ob mice that are models for type 2 diabetes and obesity respectively (47). Though there may be several possible causes behind such persistent activation of S6K1, downregulation of the negative feedback is also a probable reason.

We have observed that the IGF-1 and Insulin induce distinct dose-dependent temporal dynamics of pAkt. This is typical of temporal encoding of information in a signaling pathway. The affinity of a ligand for its receptor, phosphorylation dynamics of the receptor and interaction of adaptor molecules with the receptor determine such encoding (48–49). Network motifs at downstream parts of a pathway act as decoders and decide the flow of a signal through a particular path (19). The negative feedback of S6K1 is working like a decoder. It allows only fast rising pAkt to induce strong phosphorylation of S6K1. Thereby it allows a signal with fast rising pAkt to pass through Akt/mTORC1/S6K1 path. Accordingly, the difference in pAkt dynamics for IGF-1 and Insulin translates into differential activation of S6K1. In our experiments, 10 nM of Insulin induced transient phosphorylation of S6K1. However, such induction was very feeble. In comparison to insulin, the same amount of IGF-1 induced transient but marked increase in phosphorylation of S6K1. Therefore, one can consider that the negative feedback of S6K1 is preferentially allowing the IGF-1 signal to pass-through Akt/mTORC1/S6K1 path.

Such preferential transfer of a signal through a particular path should also modulate the effect of an external signal on a cell. S6K1 is involved in the control of cell survival and proliferation. Therefore, activation of this pathway should increase cell count. We observed that treatment with IGF-1 increased cell number dose-dependently. However, Insulin failed to induce any considerable change in cell number. Correlation analysis showed that the fold change in cell number has a reasonable correlation with the amplitude of pS6K1.

Multiple pathways and a large number of molecules control cell proliferation and survival. Therefore, the net effect of Insulin and IGF-1 signaling on cell proliferation should not be explained in terms of the activity of S6K1 only. Even then, the observed correlation between phosphorylation of S6K1 and change in cell number reiterates the critical role of signaling through Akt/mTORC1/S6K1 pathway in cell proliferation and survival. A large-scale study, involving multiple downstream branches of PI3K/Akt pathway, may help to gain a complete understanding of the differential effect of IGF-1 and Insulin on cell proliferation and survival.

## Methods

### Cell lines and culture conditions

Human Breast Cancer cell line MCF-7 was obtained from National Center for Cell Sciences, India and was cultured in Dulbecco’s modified eagle’s medium (DMEM, Himedia) supplemented with 10% fetal bovine serum (Gibco) at 37ºC in a humidified incubator with 5% CO_2_.

### Treatment of cells and Western blots

MCF-7 cells were serum starved for 16 h and treated with different doses of recombinant human IGF-1 (Gibco) and Insulin (Himedia) for different durations as mentioned in the results section. Whenever required, cells were treated with various pathway inhibitors: 10 μM of LY294002 (Sigma), 10 μM U0126 (Sigma), 10 μM PF-4708671 (Sigma), and 0.1 μM Rapamycin (Sigma).

After treatments, cells were lysed in RIPA buffer. Total protein content of each lysate was estimated by Lowry’s Method (50). An equal amount of samples were resolved by SDS-PAGE and transferred to PVDF membrane by electrotransfer. The membrane was blocked and incubated, overnight, with appropriate primary antibody at 4 ^o^C. Subsequently, blots were probed with HRP-labeled secondary antibody. List of antibodies and dilutions used are given in Supplementary Text. Blots were developed using chemiluminescence (SuperSignal West Dura kit, ThermoFisher Scientific) and imaged using gel documentation system. Densitometry of the images was performed using Image J (51). In each blot, lysate of cells treated with 10% serum was used as a sample. The densitometry readings of other samples in a blot were normalized by the reading of the serum treated sample in that blot. This allowed us to compare data from different independent blots. This normalized quantitative data has been used for further studies.

### Estimation of change in cell number

MCF-7 cells in 96-well tissue culture plates were treated with different doses of IGF-1 and Insulin for 48 hr in serum-free media. Subsequently, change in cell number was measured by a fluorescent DNA-binding dye-based method (52). In brief, cells were washed with PBS, treated with chilled methanol, followed by washing and staining with Hoechst 33342 (Sigma). Excess dye was removed by washing, and cells were treated with denatured ethanol. Fluorescence intensity of each sample was measured using a micro-plate reader (λ_exc_ = 365 nm, λ_em_ = 460 nm). Fold change in cell number was estimated by comparing fluorescence intensity of IGF-1/Insulin-treated samples with untreated cells. For cell number in the range of 5000 – 50000 cells/well, this method shows linearity between fluorescence intensity and cell number (data not shown).

### Mathematical modeling and analysis

A mathematical model, using a system of ordinary differential equations, was created for the negative feedback motif of S6K1. The equations are based on Michaelis–Menten equation. The time courses of phospho-Akt and phospho-S6K1 are input and output of this model, respectively. Parameters of the model were estimated, from experimental data, using MATLAB-based Data2Dynamics (53). Details of methods used, the differential equations, and estimated parameters are given in the supplementary text. The model was simulated and analyzed using MATLAB 2015b and JSim (54). The model in MATLAB, JSim, and SBML formats are provided as supplementary files.

## Acknowledgment

We thank Department of Biotechnology, Government of India, for financial support through Project No. BT/PR13560/COE/34/44/2015.

## Author contributions

P. D.: Designed, performed experiments, and analyzed data. V. D.: Developed and analyzed the mathematical model. B. B.: Designed the study, supervised the work, analyzed data, and written the manuscript.

## Supplementary Information

Details of reagents used, details of the mathematical model, additional data, the model in MATLAB, SBML and JSim format are provided as supplementary materials.

